# DoRIAT: A Bayesian framework for interpreting and annotating docking runs

**DOI:** 10.1101/2024.12.02.626325

**Authors:** Christos Maniatis, Zahra Ouaray, Chengkai Xiao, Thomas P.E. Dixon, Charlie Naylor, James Snowden, Michelle Teng, Jacob Hurst

## Abstract

The advent of sequence-to-structure deeplearning models have transformed protein engineering landscape by providing an accurate and cost effective way to determine crystal structures. Despite their accuracy, deep-learning predictions tend to give limited insights around protein dynamics. To improve conformation exploration we have developed a machine learning pipeline that combines deep-learning predictions with molecular docking. In this report, we propose **Do**cking **R**un **I**ntepretation and **A**nnotation **T**ool (DoRIAT). In contrast to frameworks that score models based on interface interactions, DoRIAT uses a set of parameters that summarize binding conformation. We use DoRIAT to score output from docking runs, identify complexes close to the native structure and create ensembles of models with similar binding conformations. Our results demonstrate that the single structural model DoRIAT selects to be the closest representation of the crystal structure lies within the top 10 of docked models, ranked by Root Mean Squared Distance(RMSD), in around 80% of cases.

## 1. Introduction

T cell-mediated immunity is a crucial aspect of our immune system, playing a vital role in defending against pathogens and cancerous cells. As T cells have the ability to scrutinise the entire proteome within a cell, by interrogating short peptides presented on the surface of cell by human leukocyte antigens (pHLA), they can constitute a very effective means to protect humans from pathogens or other foreign substances (Coles et al., 2020). Characterising the molecular factors governing interactions between T cell receptor (TCR) and pHLA could shed light on T cells’ ability to discriminate against antigens (Coles et al., 2020) and pave the way for developing novel T-cell therapies targeting cancer and autoimmune diseases. An example of a complete TCR-pHLA complex from the public domain, determined by X-ray crystallography (Maveyraud & Mourey, 2020), can be found in Fig. 7.

Resolving protein crystals has a long and interesting history dating back to the early 1930’s (Bernal & Crowfoot, 1934; Clark & Corrigan, 1932; Kendrew et al., 1960; HODGKIN, 1949). However, the idea that the knowledge of a protein structure could aid in the design of specific ligands, appeared a few years after the launch of the Protein Data Bank, in 1971 (Beddel et al., 1976; Bernstein et al., 1978; Maveyraud & Mourey, 2020). Technological improvements like crystallisation automates, brighter synchrotron X-ray sources, faster detectors, automated structure solution and refinement pipelines have significantly improved time (Dauter & Wlodawer, 2016; Maveyraud & Mourey, 2020), making structural resolution an integral part of pre-clinical target-based drug design. Structure-based design has now delivered drugs for a number of important diseases, including cancer, HIV, glaucoma and hypertension (Souers et al., 2013; Davis et al., 2008). However, end-to-end protein crystal structure determination remains an expensive and laborious process.

The advent of deep-learning based sequence-to-structure models like AlphaFold (Jumper et al., 2021) have transformed the landscape for *insilico* protein engineering. The initial versions of AlphaFold did not accurately model multichain complexes, however this has been addressed by a later release (Josh et al., 2024). Although modern sequence-to-structure deep-learning models can generate highly accurate structural predictions, they frequently display a significant degree of structural homogeneity. Therefore, a crucial knowledge gap persists with regards to the incorporation of molecular dynamics and flexibility, as highlighted in (Josh et al., 2024). A single structural model gives a static picture of a system which in reality is in a constant state of flux. An ensemble-based approach can overcome this problem and more comprehensively describe the preserved and transient interactions. Our approach has been to model the various TCR-pHLA complex components with sequence-to-structure models and bring them together with traditional docking algorithms, similar to (Giulini et al., 2023), in a pipeline named MLDock.

More precisely, we model alpha and beta chains of the TCR and the HLA using TCRModel2 (Yin et al., 2023), an AlphaFold derivative. Then HADDOCK3.0 (Dominguez et al., 2003), a physics based docking platform, brings together the TCR, peptide and HLA like similar methods found in literature (Cross et al., 2009; Halperin et al., 2002; Camacho & Vajda, 2002; Smith & Sternberg, 2002). The docking process involves two core steps. During the first step different ligand conformations in the active site of protein are sampled (Halperin et al., 2002). To reduce computational burden, one protein is fixed in space and the second is rotated and translated around the first one (Dominguez et al., 2003). Subsequently, correct docking poses are delineated from incorrect ones with a force-field based scoring function. A summary of the described process is shown in Fig. 6. Each docking run generates hundreds of possible protein conformations which are analyzed to suggest mutations for discovery and engineering of TCRs to create cancer therapeutics (Shafer et al., 2022).

In this report, we present **D**ocking **R**un **I**nterpretation and **A**nnotation **T**ool (DoRIAT) a method for interpreting and annotating the output of physics based docking programs like (Chen &Weng, 2002; Trott & Olson, 2010; Dominguez et al., 2003). DoRIAT is a Gaussian Process (GP) based regression model which associates root mean squared distance (RMSD) between TCR-pHLA docked model and the crystal structure (xtal) using geometric parameters derived from the TCR-pHLA complex. We use DoRIAT to give a favourable scores to conformations which are more likely to initiate immune response (i.e. canonical binders), identify complexes close to the resolved crystal structure and identify similar docking poses to create an ensemble of models with similar binding conformations. This enables a number of downstream use cases that are of deep interest for *insilico* guided protein engineering.

## 2. Methods

DoRIAT^1^ implements a Gaussian Process regressor to score MLDock models from six binding mode parameters, three angles and three displacements.

Consider *U* TCRs for which MLDock can give us *M* models. For each TCR *u* ∈ (1, …, *U*) and model *m* ∈ (1, …, *M*), **x***_um_* describing the binding mode parameters and is given by:

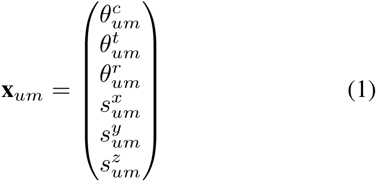

A detailed description as to how these geometric parameters are obtained can be found in Sec. A.4. The associated response **y***_um_* describing the docked model’s distance from native structure is modelled as a noisy observation of the function evaluated at **x***_um_*, f*_um_* = f (**x***_um_*) through likelihood:

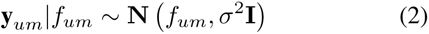

P (**y***_um_*|*f_um_*) models the data distribution at **x***_um_*and the function *f*.

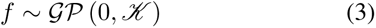

where *K* is a positive semi-definite function that determines the covariance of *f* at locations x_1_ and x_2_ and reflects our beliefs regarding the behaviour of *f*. In this work we chose the Matérn kernel described by the following formula.

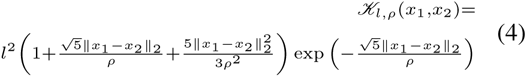

where *l* is the amplitude controlling the marginal variance of the function and ρ is the length-scale controlling spatial variability.

### 2.1. Objective function for selecting models

Our model selection strategy is a variation of µ*_RMSD_* minimisation which at the same time drastically reduces search space. More precisely, we select the best model is by:

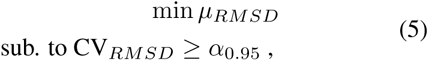

where α_0.95_ is the 95^th^ percentage quantile of CV*_RMSD_* in the docking run. Minimizing the µ*_RMSD_* is there to give the best model and CV*_RMSD_* is a constrain used to reduce the search space while resolving situations where multiple models present very similar µ*_RMSD_*. More information can be found in Sec. A.8.

This objective can be problematic when multiple models with identical µ*_RMSD_* present a coefficient of variation above threshold. In practice, we have never encountered an example like that.

### 2.2. Identifying models with similar docking poses using GP covariance

The covariance function of Eq. 3 not only encodes assumptions about the function f, but also dictates the similarity (Rasmussen & Williams, 2018) between docking poses.

We use that property to build a graph *G* (V, E) of vertices (V) and edges (E) representing binding mode parameters and connected poses respectively. The connection between poses is dictated by partial correlation, which measures dependence while accounting for effects of confounding conformations.

Let *f* (**X**) = {f*_um_* ∀u ∈ (1, …, *U*) and m ∈ (1, …, M)} be the set of RMSD predicted for *U* TCRs and M models, where *f* described in Eq. 3, **X** = x*_um_* {x*_um_* ∀u ∈(1, …, U) and *m* ∈ (1, …, *M*). Without loosing generality we chose TCR with index 1 and two conformations *m*_1_ and *m*_3_. Their predicted RMSDs are given by *f*_1,*m*1_ and _1,*m*3_ respectively. The partial correlation between variables *f*_1,*m*1_ and *f*_1,*m*3_ is a measure of their conditional association, given the remaining elements 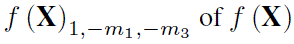 and it is defined as:

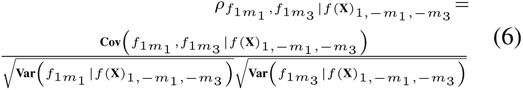

For the purpose of this project, we assume that any two models with binding mode parameters whose partial correlation is higher than 0.05 are related. This lays the foundation for choosing similar models and building ensembles around conformations of interest.

### 2.3. Contact Maps

A contact map is a two-dimensional representation used to visualize amino acid interactions between proteins of interest. Each residue in the protein complex is assigned a position on the map and colours are used to indicate interactions. We consider two residues to interact if their C*_A_* atoms are at most 8 Å apart. We use a binary representation to indicate amino acid interactions, with residues meeting our definition being labelled with 1 and 0 otherwise.

In Sec. 3 contact maps are used to asses different ensembles. We separately consider the TCR-HLA and TCR-Peptide contacts as they can give us some idea about interactions that drive affinity and specificity respectively. The contact maps of Fig. 4, 5 are created by averaging across individual contact maps of models in each ensemble.

## 3. Results

DoRIAT is a tool for the exploratory analysis of TCR-pHLA structures produced by docking simulations. It aims to establish a robust docking model scoring system from six binding mode parameters characterising the relative position of the TCR to the pHLA, briefly described in Tab. 1 and more detailed in Sec. A.4. In turn, this is used for a set of downstream tasks such as identify and rank near-native conformations or detect similar conformations. In this paper we will focus on docking outcomes from MLDock, our in-house built docking pipeline, however similar approach could be used to understand simulated conformation from any docking engine.

**Table 1.** Nomenclature. Table briefly describing parameters used to summarize TCR-pHLA complexes. Binding mode parameters are bolded. A detailed description of how these parameters are computed can be found in Sec. A.4

|  |  |
| --- | --- |
| $v_{rot}$ | Alpha and beta chain rotation symmetry vector pointing towards the HLA. |
| $v_{SB}$ | Vector between the disulphide bonds' centroids in alpha and beta chain pointing towards beta chain. |
| $p_{TCR}$ | TCR's centre of mass. |
| $p_{MHC}$ | MHC's centre of mass. |
| $v_{MHC_1}, v_{MHC_2}$ | Vectors defining the TCR-pHLA interface plane. |
| $n_{MHC}$ | Vector perpendicular to $v_{MHC_1}$ and $v_{MHC_2}$ pointing towards TCR. |
| $\theta^c$ | Angle between $v_{MHC_1}$ and $v_{SB}$. |
| $\theta^t$ | Angle between $v_{MHC_1}$ and $v_{rot}$. |
| $\theta^r$ | Angle between $v_{MHC_2}$ and $v_{rot}$. |
| $s^x$ | $v_{MHC_1}$ direction displacement of $p_{TCR}$ from $p_{MHC}$ in the interface plane. |
| $s^y$ | $v_{MHC_1}$ direction displacement of $p_{TCR}$ from $p_{MHC}$ in the interface plane. |
| $s^z$ | Distance of $p_{TCR}$ projected TCR-pHLA interface plane. |

TCRs can bind in multiple conformations to pHLA. This is also highlighted by the hundreds of models that rank well according to HADDOCK’s scoring function, which serves as a proxy for affinity. When TCR-pHLA binding exceeds geometric tolerances, important cell surface interactions do not take place and signaling is disrupted (Adams et al., 2011). We refer to docking geometries that fail to induce activation as non-canonical and we have actively tried to remove them in post-docking analysis. Our first approach for identifying and eliminating non-canonical poses was to derive thresholds from limited publicly available X-ray crystallography structures which can be found in Tab. 2. As the structures used to estimate these ranges presented significant levels of similarity and were different from the systems we were internally interested, their use created challenges in our engineering effort. To overcome the problem of setting geometric limitations, we redefine the problem as a prediction of distance to the crystal structure, where signaling is more likely. We developed DoRIAT, a GP regressor based scoring function that takes into account combinations of binding mode parameters to identify models close to the native structure, schematically described in Fig 6.

**Table 2.** Table summarizing the cut-off thresholds for binding mode parameters.

| | $\theta^c$ | $\theta^t$ | $\theta^r$ | $s^x$ | $s^y$ | $s^z$ |
| --- | --- | --- | --- | --- | --- | --- |
| min | 19 | -25 | -30 | -17 | -7 | 20 |
| max | 75 | 25 | 25 | 12 | 10 | 30 |

Briefly, for each TCR u, DoRIAT associates RMSD distance **y***_um_* between C*_A_* atoms of docked model m with its corresponding crystal structure with TCR-pHLA binding mode parameters **x***_um_* using a GP f. Our kernel of choice for the GP is the Matérn with 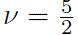 with amplitude l and lengthscale ρ. As the RMSD data are initially scaled between [0, 1] and subsequently inverse-transformed by the cumulative cdf of a standard normal, observation noise model is a normal distribution with mean the output of the GP *f* and variance σ^2^. More details can be found in Sec. A.4. DoRIAT then constructs a constrained optimization problem involving the predicted RMSD and its corresponding coefficient of variation (CV) to select a model close to the native structure. DoRIAT’s covariance matrix is used to identify similar docking conformations allowing for the creation of ensembles around points of interest.

### 3.1. DoRIAT accurately scores output models of docking run

DoRIAT is trained on docking output of 43 TCRs and subsequently tested on 15 TCRs sourced from STCRDab (Leem et al., 2018). For each TCR the docking platform generates 600 possible conformations of the TCR-pHLA complex. Both training and testing data comprise of a mixture of complexes of a TCR bound to viral or cancer peptides presented by HLA-A^∗^02. For testing we also include proprietary data.

Fig. 1 summarizes the scoring results for docked conformations of HIV (5NMG), viral (5ISZ) and cancer (2PYE) related systems from the test set. DoRIAT accurately predicts docked models far from the native structure, where binding mode parameters range outside the canonical range and can be identified with a thresholding approach as indicated by the orange and blue color-coding. DoRIAT struggles with models closer to the native structure, as indicated by larger deviations in Fig. 1c and 1b. In these examples, some of the docked models present binding mode parameters that are on the edge of what the threshold-based approach would deem canonical. However, there are several examples like 5NMG (Fig. 1a), where such edge cases are absent and DoRIAT is very accurate.

**Figure 1.**
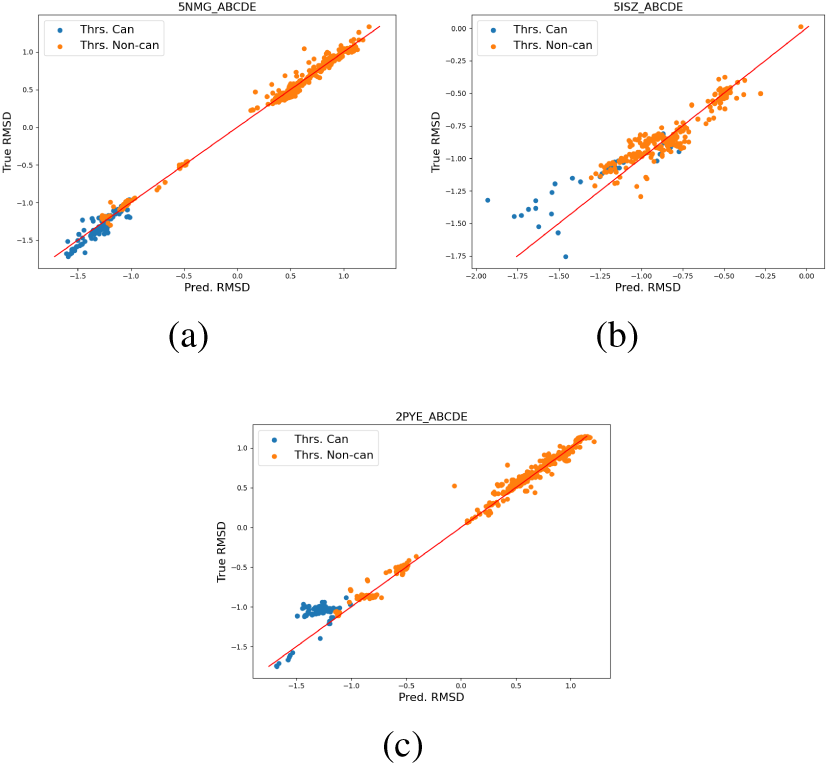
Measured RMSD as a function of predicted RMSD for three TCRs in the test set. Measured RMSD as a function of predicted RMSD for crystal structures 5NMG (Fig. 1a), 5ISZ (Fig. 1b) and 2PYE (Fig. 1c). Each dot represents a docked model and is color-coded using cut-off thresholds of Tab. 2. The red line corresponds to perfect match between predicted and measured RMSD.

**Figure 2.**
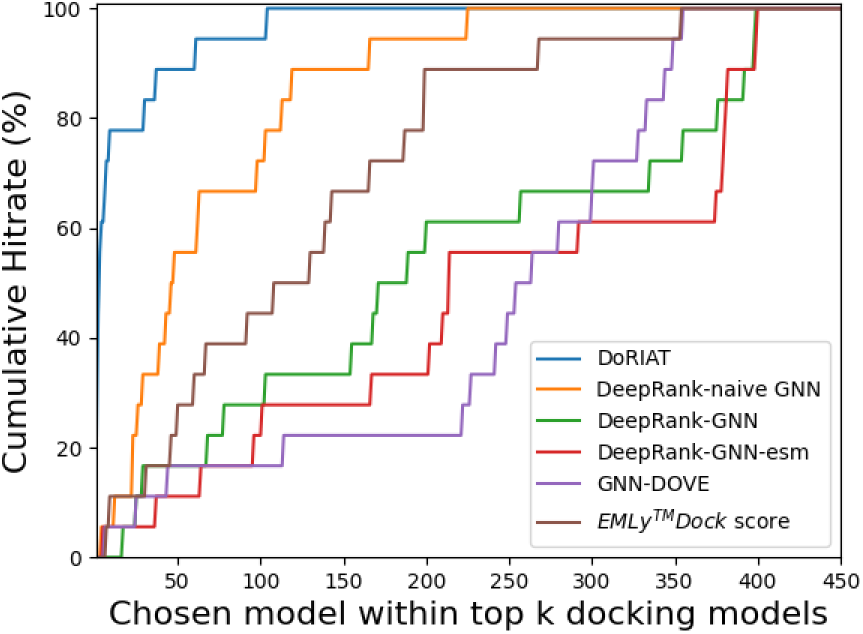
Cumulative hitrate across compared methods. Graph summarizing cumulative hitrate for DoRIAT (blue), naive GNN (orange), DeepRank-GNN (green), DeepRank-GNN-esm(red) and MLDock (purple) across unseen complexes. Each method chooses one docked model that it believes is closest to the crystal structure. These picks are then ranked using the measured RMSD from crystal structure. We set the lowest ranking to 450 as all methods reach 100% cumulative rate before this position.

For TCR-pHLA systems like 2PYE and 5ISZ, where DoRIAT’s predictions differ from the ground truth, the native complex parameters are listed in the first two lines of Tab 3. For 2PYE and 5ISZ, models like 2PYE m1 and 5ISZ m1 have parameters θ*^t^* and θ*^c^* close to thresholds presented in Tab 2 and highlight challenges for threshold-based method. In absence of ground truth the presented models could be considered plausible TCR-pHLA complexes, even though their backbone distance from their corresponding native structures is greater than 11.5 Å. While DoRIAT underestimates the models’ distance from native structure, it is still able to suggest that all considered models are more than 10 Å away from crystal structure, unlike the threshold based approach which would incorrectly retain them.

DoRIAT was also applied to docking simulations of unpublished TCR-pHLA complexes, which are more heterogeneous than public data and intended for cancer therapeutics. Their binding mode parameters can be found in rows 3 and 4 of Tab. 3. A direct consequence of the inherent complexity is to have conformations with parameters within range, whose backbone is far from crystal structure causing DoRIAT to struggle with accurate RMSD prediction. TCR A’s native structure has (θ*^c^*, θ*^r^*) angles of (40.56^◦^, 10.76^◦^), while TCR-pHLA complex A m1 and TCR-pHLA complex A m2 bind at (22.75^◦^, 13.67^◦^) and (70.17^◦^, 16.12^◦^). For these examples, the discrepancy between DoRIAT’s prediction and actual RMSD would be 5 Å and 3 Å respectively. However, these models would not affect downstream analysis as they would easily be flagged for removal due to their high predicted RMSD (19 Å and 7 Å respectively). TCR B’s listed in Tab. 3 (row 4) is one of the most difficult TCRs to make predictions. TCR-pHLB complex B m1 and TCR-pHLA complex B m2 are part of a bigger trend presented in Fig. 3b, where DoRIAT underestimates the RMSD from crystal structure as binding mode parameters are far from native structure, but within range.

**Figure 3.**
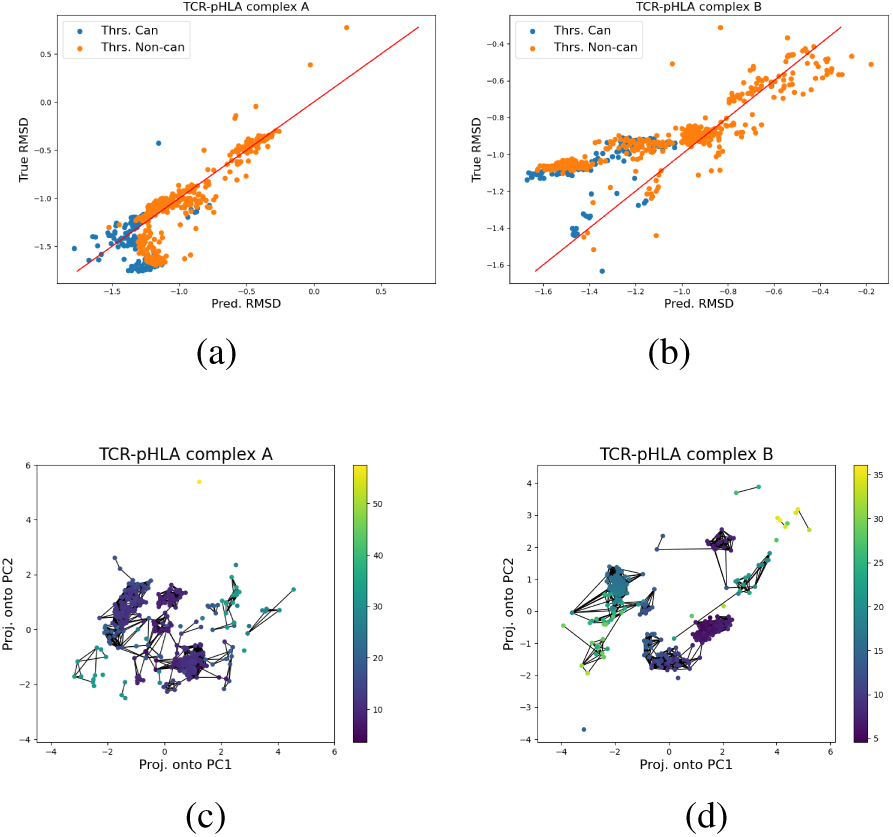
Interpretation of docking runs for internal data. Figures summarizing post-docking analysis for TCR A and TCR B. (3a,3b) Scatter plots of measured RMSD as a function of predicted RMSD for TCRs that DoRIAT performs well (TCR A) and struggles (TCR B) respectively. Each dot represents a docked model and is color-coded using cut-off thresholds of Tab. 2. The red line represents a perfect match between predicted and measured RMSD. (3c,3d) PCA embedding of binding mode parameters for TCR A and TCR B, color-coded by predicted RMSD. Each dot represents a docked model. Each dot corresponds to a docked model, with edges connecting conformations that share similar binding mode parameters, as determined by partial correlation from GP covariance.

**Figure 4.**
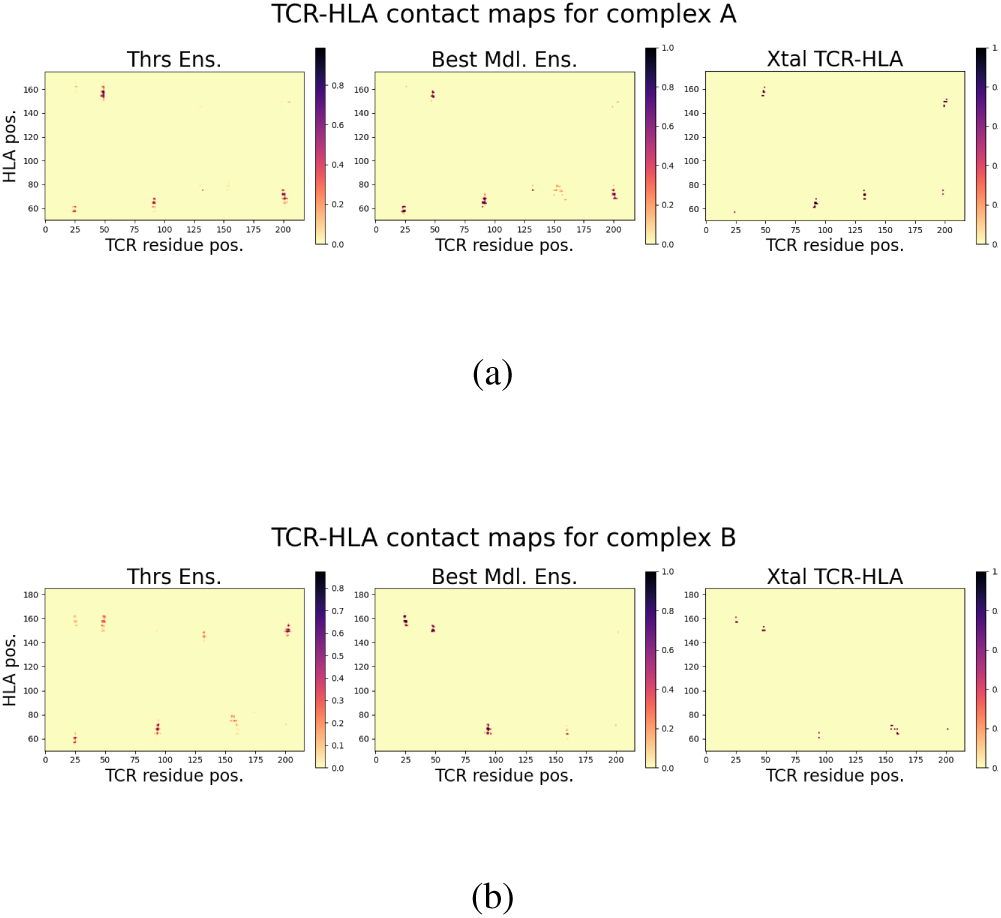
Contact maps between TCR-HLA. (8 Å) **for different ensembles for complexes A and B.** Summary of contact maps between the TCR-HLA for complexes A (Fig. 4a) and B (Fig. 4b) for ensembles created using canonical models based on thresholds and similar models around prediction made with Eq. 5. Here a contact is defined if two amino acids have a distance of less than 8 Å. Each contact map is created by averaging contact maps of models in each ensemble. Positions on the contact map are colored based on the average contacts between TCR and HLA. For comparison we include the contact maps observed in the crystal structure.

**Table 3.** Binding mode parameters for section’s examined examples. Summary of the binding poses for native complexes (first four lines) and docking outcomes with canonical parameters that represent structures distant from the ground truth (remaining six lines).

| | $\theta^c$ | $\theta^t$ | $\theta^r$ | $s^x$ | $s^y$ | $s^z$ |
| --- | --- | --- | --- | --- | --- | --- |
| 2PYE | 66.73 | 8.21 | -17.03 | 4.39 | -4.96 | 24.74 |
| 5ISZ | 56.33 | 2.66 | -7.34 | 3.34 | 2.71 | 24.53 |
| TCR A | 40.56 | 6.40 | 10.76 | 0.72 | -5.18 | 23.06 |
| TCR B | 65.67 | -3.74 | -8.53 | 2.87 | -0.29 | 25.29 |
| 2PYE.m1 | 41.31 | 23.97 | 8.88 | 4.66 | 1.12 | 22.71 |
| 5ISZ.m1 | 21.75 | 19.46 | 5.81 | 11.48 | -1.04 | 22.14 |
| TCR-pHLA complex A.m1 | 22.75 | 22.49 | -13.67 | 9.75 | -6.61 | 23.08 |
| TCR-pHLA complex A.m2 | 70.17 | -4.87 | -16.12 | 6.02 | -6.93 | 23.59 |
| TCR-pHLA complex B.m1 | 21.55 | 7.06 | 9.32 | 6.69 | -0.53 | 24.08 |
| TCR-pHLA complex B.m2 | 45.56 | -18.38 | -6.85 | -2.42 | -6.46 | 24.85 |

By jointly evaluating binding mode parameters, DoRIAT is more effective than the threshold-based method, offering higher control over the granularity of the post-docking analysis. However, DoRIAT’s training on limited data creates unique challenges in subsequent tasks like finding a model close to the native structure. In following section, we out-line an approach that leverages the model to address this challenge.

### 3.2. DoRIAT outperforms state of the art in identifying models close to the native structure

By benchmarking MLDock on public and internal TCR-pHLA complexes, we have identified that the median RMSD between the best simulated pose and the crystal structure is 3.1 Å. This can become particularly useful when structurally assessing candidates from our screening for therapeutics pipeline where access to a crystal structure is not available. Having a method that can suggest a docked models close to the native structure allows to cost-effectively evaluate if a candidate has engineering potential.

The comparison between different model selection approaches is performed using 18 TCR-pHLA complexes, 15 from DoRIAT’s test set and 3 internal which include TCR-pHLA complex A and TCR-pHLA complex B. Internal structures are cancer related, while public structures contain a mixture of viral and cancer. All structures are bound to HLA-A^∗^02.

To evaluate the effectiveness of different methods in identifying models close to the native structure, we consider, alongside DoRIAT, MLDock score (HADDOCK score in water refinement), three DeepRank variations and GNN-DOVE. HADDOCK score is a weighted sum of electrostatic, van der Waals, desolvation and distance restraints energy functions to discriminate native looking structures from the rest. Hence, the lower the score the closer to the native structure should be. For DeepRank we train a naive Graph Neural Network (naive-GNN) and DeepRank-GNN (Réau et al., 2023) to predict RMSD using the code provided in DeepRank2. For DeepRank-GNN-esm (Xu & Bonvin, 2024) and GNN-DOVE, we use the pre-trained models provided in DeepRank-GNN-esm and GNN-DOVE to score models.

All models are set to their default hyper-parameters and are trained on the same data split as DoRIAT using Adam optimizer with 5 step patience early stopping policy for a maximum of 20 steps. For the naive GNN and DeepRank-GNN the computationally intensive PSSM step is skipped.

For each of the compared methods, one docked conformation is selected based on each method’s scoring approach. For DoRIAT the model is selected such that it solves Eq. 5. Model selection on DeepRank variations naive-GNN and DeepRank-GNN is performed based on predicted RMSD. For DeepRank-GNN-esm and GNN-DOVE selection is based on fraction of native contacts and the probability that the docking decoy has a CAPRI acceptable quality respectively. For MLDock selection is based on lowest HADDOCK score. These predictions are then ranked based on their RMSD from crystal structure. Subsequently, we define the hitrate as a binary variable specific to each method which is 1 for the selected position and 0 otherwise.

Fig. 2 summarizes the cumulative hitrate as a function of ranking. The structural model that DoRIAT selects to be the closest representation of the crystal structure, out-performs compared methods by being within top 10 of docked models, ranked by RMSD, 78% of times (14 out of 18) with 4 times identifying the best model. In comparison, Naive GNN and MLDock identify models within the top 70 in 60% and 40% of cases respectively, while DeepRank-GNN, DeepRank-GNN-esm, and GNN-DOVE achieve only 22%, 17%, and 17%. More results can be found in Tab. 5. Since the docking engine effectively explores the space of conformations and creates plausible interfaces, predicting canonical models from local interactions becomes challenging. Hence, using binding mode parameters gives DoRIAT significant benefit over alternatives. DeepRank’s performance further degrades due to the lack of PSSM, which was ignored as it would make unfeasible the assessment of large TCR batchess. The DeepRank-GNN-esm and GNN-DOVE have been pre-trained on antibody-antigen docking results, which differ from our TCR-pHLA systems, therefore performance is not optimal. However, if we were to retrain or fine-tune them on DoRIAT’s training data we would expect them to face some of the challenges encountered by the other DeepRank variations. MLDock’s scoring function is the third highest performer with the exception of TCR-pHLA complex A where it performs better even compared to DoRIAT.

To further assess the quality of selections, we consider DockQ score (Basu & Wallner, 2016) which incorporates additional measures like F_nat_, LRMS and iRMS standardized by CAPRI (Lensink & Wodak, 2013). Tab. 4 and Tab. 6 summarize the DockQ scores for model selection in benchmarking across test set and internal TCRs. DoRIAT presents superior performance with all but one selection having at least acceptable quality and an average DockQ score of 0.42. The second best performer is the naive-GNN DeepRank variation with 5 out of 18 selected models having acceptable quality and an average DockQ score of 0.24. The rest DeepRank based models, MLDock and GNN-DOVE have 7 or fewer acceptable models and an average score of less than 0.20. Furthermore, some of the selections made by DeepRank variations, GNN-DOVE and MLDock are not guaranteed to be within canonical range.

**Table 4.**
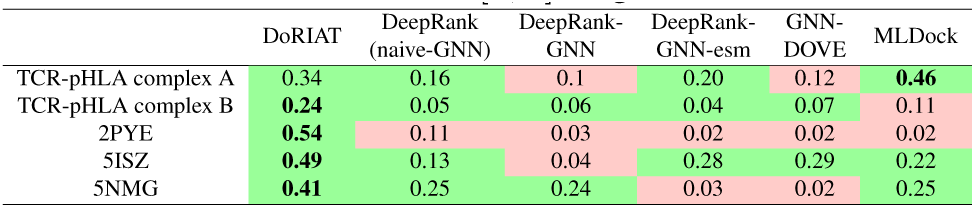
Assessment of model selections using DockQ score for test set and internal TCRs. (Part1) Best perfroming selection is marked with bold. Cells are color-coded to indicate whether selec-tion is canonical (green) or not (red) according to the thresholds of Tab. 2. DockQ scores are in the [0, 1] range.

|  | DoRIAT | DeepRank<br>(naive-GNN) | DeepRank-GNN | DeepRank-GNN-esm | GNN-DOVE | MLDock |
| --- | --- | --- | --- | --- | --- | --- |
| TCR-pHLA complex A | 0.34 | 0.16 | 0.1 | 0.20 | 0.12 | **0.46** |
| TCR-pHLA complex B | **0.24** | 0.05 | 0.06 | 0.04 | 0.07 | 0.11 |
| 2PYE | **0.54** | 0.11 | 0.03 | 0.02 | 0.02 | 0.02 |
| 5ISZ | **0.49** | 0.13 | 0.04 | 0.28 | 0.29 | 0.22 |
| 5NMG | **0.41** | 0.25 | 0.24 | 0.03 | 0.02 | 0.25 |

In the left-out set, DoRIAT is sometimes outperformed by other methods based on DockQ scores, such as in TCR-pHLA complexes A, TCR-pHLA complex C, 3QFJ and 4MNQ. For the first two, MLDock’s selections are slightly better. However, TCR-pHLA complex C has x-axis and y-axis displacements (s*^x^*, s*^y^*) of (13.1 Å, 9.4 Å), which is outside of canonical range. For 3QFJ and 4MNQ naive-GNN method performs best. In all these examples, the main reason other methods get better DockQ scores is attributed to better interface quality. This is expected as DoRIAT focuses on the global geometric properties of the complex. Even in these cases, DoRIAT’s selections fall into the same CAPRI quality class with top performers. Overall, DoRIAT exhibits greater consistency in the quality of selections.

In presence of larger training dataset, we would expect the disparity between DoRIAT, naive-GNN and DeepRank-GNN performance on post-docking analysis to close. However, DoRIAT alternatives would struggle as modern docking engines are effective at creating plausible interfaces even for non-canonical conformations. Hence, in this setting, binding mode parameters provide a robust alternative for building machine learning models without the need for massive amounts of data.

### 3.3. DoRIAT covariance can help resolve the post-docking landscape

Fig. 2 demonstrates that DoRIAT often selects models close to the crystal structure. Beyond that, DoRIAT can also discriminate between poses, giving clear distinction between disconnected conformations and those with similar global geometry, enabling the creation of model ensembles around points of interest. This expands the limited perspective offered by crystal structures or individual models. In this section, we briefly overview how DoRIAT’s covariance is used to create ensembles around points of interest using two complexes, TCR-pHLA complex A and TCR-pHLA complex B.

Fig. 3c and 3d show PCA of binding mode parameters for complexes A and B, with edges connecting similar conformations. In both examples docking poses cluster around distinct canonical and non-canonical poses. For complex A conformations within canonical range have θ*^c^* of about 35^◦^ and 45^◦^, while for complex B canonical conformation cluster around θ*^c^* of about 24^◦^ and 65^◦^. In absence of a crystal, an ensemble could be created either by considering all canonical models or by picking a model of interest and building the ensemble around it.

To illustrate how covariance information could aid in a more targeted analysis, we create two ensembles of models and compare their average contact maps to those of the crystal structure. The first ensemble includes models which are deemed canonical based on Tab. 2 thresholds. The second uses the optimal model from Eq. 5 along with similar models identified via partial correlation coefficient. Ensemble quality is assessed by comparing TCR-HLA and TCR-peptide contact maps to crystal contact maps using mean structural similarity index (SSIM) (Wang et al., 2004). SSIM is better suited for this task, compared to other distance metrics such as mean squared error, as it accounts for contact position. Proximity to crystal contacts is more tolerable, reflecting protein flexibility, while distant deviations are penalized more heavily as they likely indicate modeling artifacts.

Fig. 4 and 5 summarize TCR-HLA and TCR-peptide contact maps for the two ensemble methods. In Fig. 4a, both the threshold and optimal model threshold introduce artificial contacts compared to the crystal structure, notably at (TCR, HLA) = (25, 52). At the same time, both ensembles identify many correct contacts, resulting in SSIM scores of 0.979 and 0.975, with threshold-based ensemble being slightly better. For the TCR-epitope contact map (Fig. 5a), the optimal model ensemble achieves an SSIM 0.815 against 0.805 for the threshold-based method, due to fewer contacts in incorrect epitope regions. Fig. 4b suggests that for complex B the ensemble around optimal model matches the crystal contacts, while the threshold-based approach presents erroneous contacts, such as (TCR, HLA) = (26, 55), (204, 150). In this case, SSIM for the TCR-HLA contact map of the threshold-based ensemble is 0.970, 0.015 lower than that of the optimal model ensemble. The optimal model ensemble, for TCR-peptide contacts, misses contacts at (TCR, Peptide) = (200, 5) and adds incorrect ones at (TCR, Peptide) = (30, 3) (Fig. 5b). The threshold-based ensemble correctly identifies contacts at (TCR, Peptide) = (200, 5), but introduces contacts in the regions (TCR, Peptide) = (30, 3), (48, 3) (Fig. 5b). SSIM for the the optimal model ensemble contact map is 0.850 against 0.843 for the threshold-based ensemble.

**Figure 5.**
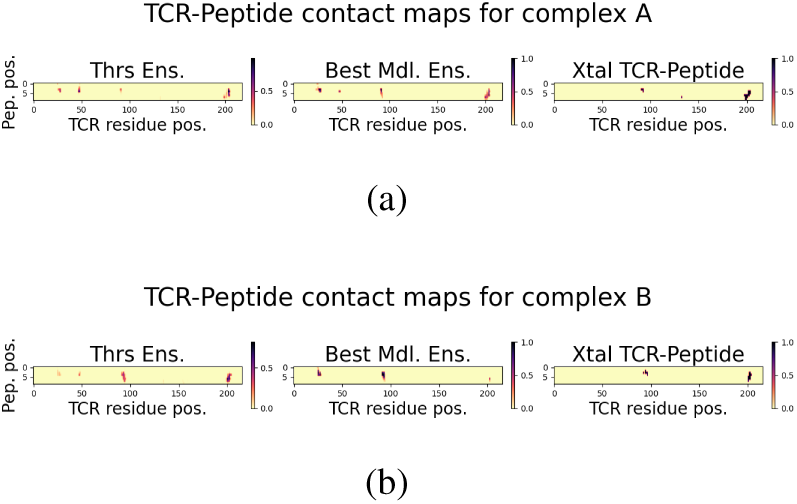
Contact maps between TCR-peptide. (8 Å) **for dif-ferent ensembles for complexes A and B.** Summary of contact maps between the TCR-peptide for complexes A (Fig. 5a) and B (Fig. 5b) for ensembles created using canonical models based on thresholds and similar models around prediction made with Eq. 5. Here a contact is defined if two amino acids have a distance of less than 8 Å. Each contact map is created by averaging contact maps of models in each ensemble. Positions on the contact map are col-ored based on the average contacts between TCR and peptide. For comparison we include the contact maps observed in the crystal structure.

In summary, our results suggest the covariance information can create ensembles with contact maps of equal quality or better quality than those derived from threshold-based methods on binding mode parameters. This enables more accurate interface analysis, improving downstream analysis.

## 4. Conclusion

The advent of sequence-to-structure models and their combination with more traditional physics based docking platforms has opened up new possibilities for *insilico* protein engineering. However, traditional docking engines introduce, among native conformations, a series of docking poses which are either impossible to activate the T cell or are far from the native structure observed in X-ray crystallography. We introduce DoRIAT, a Gaussian process regressor which uses binding mode parameters to interpret docking output. By considering combinations of binding mode parameters instead of a threshold-based approach, DoRIAT can reliably generalize on TCR-pHLA complexes of interest.

DoRIAT has been benchmarked in a set of tasks vital for *insilico* protein engineering. One of DoRIAT’s standout features is its ability to identify models close to the crystal structure. In contrast to compared methods which heavily rely on interface contacts, DoRIAT’s strategy of assessing docked conformations using binding mode parameters allows for accurate and scalable identification of models near to the crystal structure. Additionally, by leveraging DoRIAT’s covariance, conformations that exhibit similar binding geometry can be selected creating an ensemble of models around points of interest. These ensembles could be used for a more thorough interface analysis of TCR-pHLA systems of interest. Overall, the described capabilities offer a cost effective solution for evaluating engineering candidates.

DoRIAT has been developed as tool for interpreting MLDock’s output. Over time it has offered important insights in the effort to engineer picomolar affinity TCRs. We believe that insights provided in this report will offer an alternative way of analyzing docking results which extends beyond TCRs. A similar method to DoRIAT could be developed to analyze antibody-antigen complexes. This is going to come with its own challenges as antibodies lack co-evolutionary constraints, but their large numbers in public repositories could help the algorithm to attain reasonable predictive power.

## Impact statement

The development and application of DoRIAT address critical challenges in molecular docking interpretation, enabling a systematic and data-driven approach to analyzing docking results. This tool has the potential to accelerate drug discovery pipelines and improving the identification of high-potential drug candidates. As a result, DoRIAT stands to impact computational chemistry, structural biology and pharmaceutical research communities.

## Lay Summary

T cell-mediated immunity is a cornerstone of the body’s defense system, protecting against infections and cancer. Understanding the molecular interactions between T cell receptors (TCRs) and pHLA molecules is key to advancing therapies for cancer and autoimmune diseases. To address this challenge, we have developed a cutting-edge platform that combines deep-learning-based sequence-to-structure models with traditional docking algorithms to analyze TCR-pHLA interactions. At the heart of this effort is DoRIAT, a tool that enables the generation and interpretation of hundreds of potential protein conformations, identifying the most promising candidates for novel T cell therapies.

DoRIAT is a tool for interpreting docking results. It scores and prioritizes conformations most likely to trigger immune responses, identifies models closely resembling experimental crystal structures, and creates ensembles of binding models for more detailed analyses. This unique approach bridges the gap between static protein modeling and the dynamic reality of molecular interactions.

By leveraging this innovative combination of AI and molecular docking, we are positioned to redefine insilico protein engineering. Our solutions offer a scalable, cost-effective way to accelerate the discovery and optimization of T cell therapies, driving us toward a future of transformative treatments for cancer and autoimmune diseases.

## A. Appendix

### A.1. Schematic representation of the pipeline

**Figure 6.**
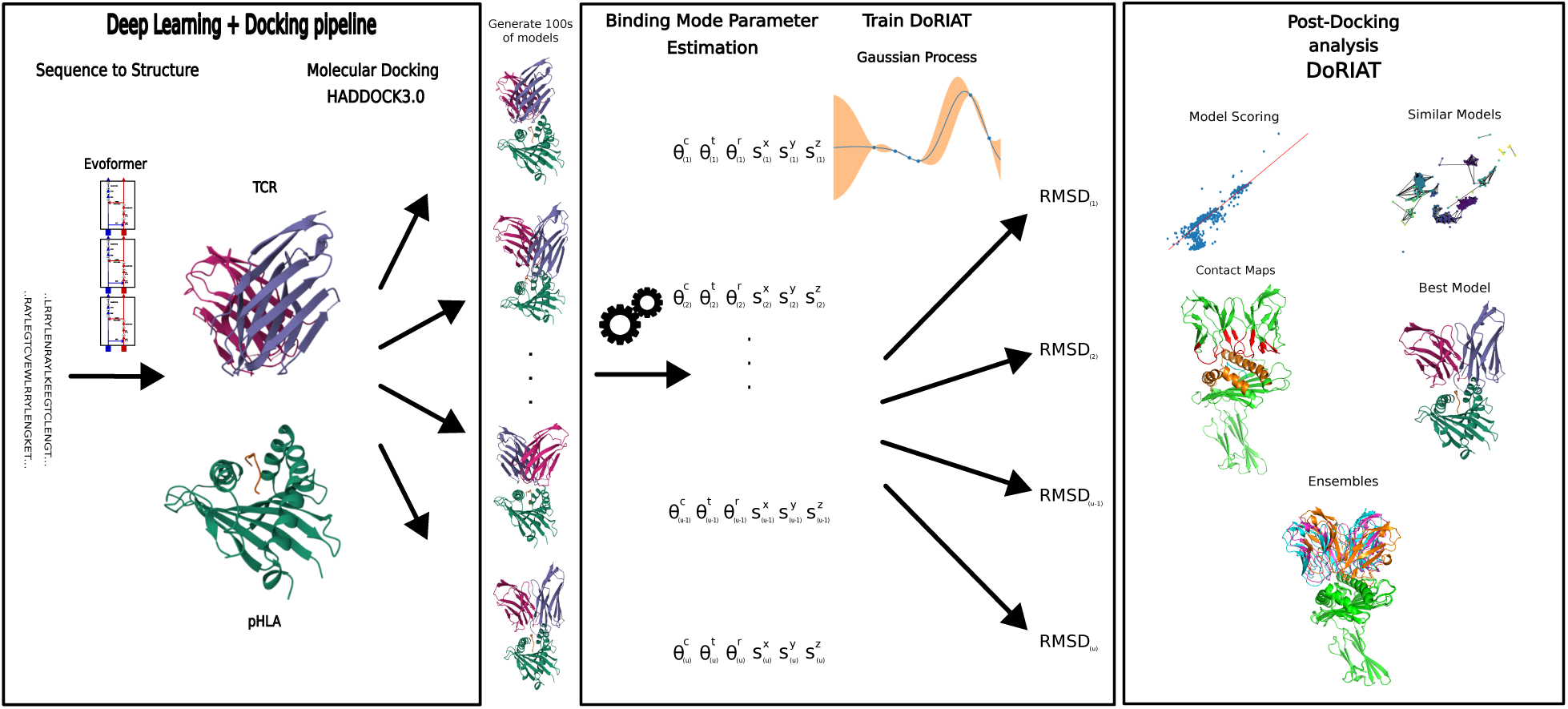
Schematic representation of MLDock and DoRIAT. In-house developed pipeline that combines deep-learning based sequence-to-structure model to generate alpha and beta chains, followed by insilico docking with HADDOCK3.0 (Dominguez et al., 2003) and DoRIAT to interpret docking outcomes and identify the best conformation.

### A.2. Related Work

The binding of proteins and other molecules is fundamental to numerous biological processes. However, predicting binding affinity from structural information alone, within reasonable time, is a challenging task, particularly when experimental data is limited or unavailable. Field practitioners have developed a plethora of approaches which are either used as part of the docking process to score conformations, or post-docking to remove some of the spurious conformations that are less likely to initiate immune response. In this section we review these two approaches explaining some of their underlying assumptions.

Docking scoring functions were developed as a proxy of protein-ligand binding affinity and can be divided into geometric matching, force-field-based, empirical and knowledge-based (Kitchen et al., 2004; Meng et al., 2012).

**Geometric matching** between protein and ligand has repeatedly reaffirmed to have a vital role in determining the geometric complex (Zielenkiewicz & Rabczenko, 1984; Kuntz et al., 1982; Halperin et al., 2002), with early docking algorithms heavily capitalising on it (Fischer et al., 1995; Norel et al., 1994a;b; 1999b;a). However, such efforts were quickly abandoned as they struggle to generalise in absence of ground truth introducing many false positives (Halperin et al., 2002).

**Force-field-based** scoring functions (Carlson & Jorgensen, 1995; Aqvist et al., 2002; Kollman, 1993) assess the binding energy by calculating the sum of electrostatics, calculated by a Coulombic formulation and van der Waals interactions described by the Lennard-Jones potential function (Meng et al., 2012). To address the slow computational speed of forcefield-based scoring functions, distant amino acid interactions are ignored. To improve upon standard force-field functions a series of extensions involving hydrogen bonds and solvation energy are included (Jackson, 2002; Moitessier et al., 2008).

In **empirical scoring** functions (Head et al., 1996; Jain, 1996; Verkhivker et al., 2000; Gehlhaar et al., 1995; Böhm, 1998), binding energy decomposes into energy components, such as hydrogen bond, ionic interaction, hydrophobic effect and binding entropy (Meng et al., 2012). The contribution of each component is estimated from a set of ligand-protein complexes with measured affinity. Many modern docking engines such as (Dominguez et al., 2003; Chen & Weng, 2002) are equipped with empirical scoring functions, accounting for a plethora of described phenomena that contribute to affinity. The main limitation of empirical scoring functions is that they might fail to generalize beyond the set of training complexes.

**Knowledge-based** scoring functions (Muegge & Martin, 1999; Mitchell et al., 1999; Ishchenko & Shakhnovich, 2002; Feher et al., 2003; Verkhivker et al., 1995) are derived by looking into the inter-atomic contact frequencies and protein-ligand distances. Their key assumption is that favorable interactions will be more frequently observed (Meng et al., 2012). Similar to empirical-based scoring approaches, knowledge-based ones often fail to generalize outside of their training data.

**Consensus scoring** (Charifson et al., 1999) combines several different scores to assess the docking conformation. It has been reported (Feher, 2006; Charifson et al., 1999; Bissantz et al., 2000; Terp et al., 2001) that consensus scoring improves prediction of bound conformations and poses (Meng et al., 2012). However, binding energies might still be inaccurate and its predictive power depends on the independence of scoring functions it is made of (Kitchen et al., 2004; Feher, 2006; Charifson et al., 1999; Bissantz et al., 2000; Terp et al., 2001).

A limitation shared across the scoring functions described above is their limited treatment of solvation energy. More rigorous approaches for estimating binding affinity would include Free Energy Perturbation (Zwanzig, 1954), Replica Exchange Free Energy Perturbation (Meng et al., 2011), Molecular Mechanics Poisson-Boltzmann solvent-accessible surface area (MM-PB/SA) (Kollman et al., 2000). However, they are computational demanding and can easily become prohibitively expensive for larger molecules(Miller et al., 2012). A work-around would be to use them for scoring post-docking.

The introduction of powerful hardware accelerators like GPUs and TPUs along with parallel file system technologies has given rise to machine learning approaches. Machine learning methods are extremely flexible and in numerous cases have proved their ability to extract patterns from millions of data points. (Wang et al., 2021) have introduced Graph Neural Network–based DOcking decoy eValuation scorE (GNN-DOVE), a method that scores docking models by looking at atom chemical properties and inter-atom distances in the TCR-pHLA interface. Similar to GNN-DOVE, DeepRank and its variations (Renaud et al., 2021; Réau et al., 2023; Xu & Bonvin, 2024) use graph neural networks (GNNs) to learn residue level features and score biomolecular complexes. Subsequently they could be used for binding affinity prediction but also to disentangle crystal artefacts from protein interactions of potential biological interest.

### A.3. Information driven docking

Information-driven docking integrates experimental or computational data to improve the prediction of biomolecular interactions, enhancing accuracy and efficiency (Dominguez et al., 2003). HADDOCK uses Ambiguous Interaction Restraints (AIRs) to handle incomplete or ambiguous data, enabling the modeling of flexible interactions (Zundert et al., 2016). Unlike unambiguous restraints, AIRs allow for multiple possible interactions between residue groups, making them ideal for uncertain data. This flexibility enables HADDOCK to explore a broader range of conformations while still being guided by the available data (Honorato et al., 2021).

To systematically explore the interaction space, we devised three distinct docking protocols, each defining active and passive residues differently. Protocol A a more permissive approach, with only a minimal number of active residues, allowing for a less constrained exploration. In contrast, protocols B and C were designed to be more targeted, by employing a larger set of active residues in the binding interface. This tiered approach enables a balanced exploration of the conformation space, combining the flexibility of Protocol A with the precision of Protocols B and C, thereby capturing both broad and specific aspects of the interaction landscape.

To train DoRIAT, we used all three protocols. For best model selection we choose the best model from subset of models docked with Protocols B and C, but the ranking was done on all three data. The ensembles were created from Protocol B and C models.

### A.4. Binding mode parameters

Binding mode parameters are essential in assessing T cell activation and subsequent immune response. DoRIAT depends on three angles and three displacements. To estimate these parameters, 2 vectors and a centre of mass for the TCR and pHLA have to be characterized respectively. Here we describe the procedure to obtain DoRIAT features and summarize the process into a pseudo-code.

Let v*_rot_* be the vector parallel to the alpha and beta chain rotation symmetry axis pointing towards the HLA, p_TCR_ be the TCR’s centre of mass, and v_SB_ the vector in the direction defined by alpha and beta chain disulphide-bridges pointing towards beta chain. Let a_alpha_ and a_beta_ be the set of amino acids’ carbon alpha (C*_A_*) atomic coordinates in alpha and beta chains respectively aligned based on the IMGT numbering (Lefranc, 1997; 1998; 1999). Let J be the indices of subsets of a_alpha_ and a_beta_ with common IMGT numbering and a_alpha_*_,J_* and a_beta_*_,J_* the relevant subsets. We consider only points with common IMGT numbering in order not to tip the centre of mass and symmetry axis towards the beta chains which is commonly longer. Let v_TCR_ be the set of midpoints between alpha and beta chain atomic C*_A_* coordinates with the same IMGT numbering.

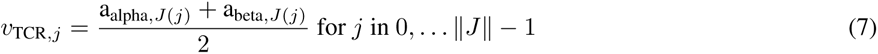

Then v*_rot_* is the normalized vector pointing towards the HLA parallel to the best fit line of v_TCR_ coordinates and p_TCR_ is the mean of v_TCR_ coordinates. v_SB_ is the normalized vector parallel to the line connecting disulphide bonds in alpha and beta chains, with the vector pointing towards beta chain.

Let v_H1_ and v_H2_ be the normalized vectors parallel to the best fit lines running through C*_A_* atoms’ coordinates in HLA helix 1 and 2 respectively. Helices 1 and 2 are defined to be the parts of HLA with IMGT numbering ranges 52-88 and 141-176 respectively. To make results consistent when estimating v_H1_ and v_H2_ we make sure the vectors point towards the last residue in the amino acid sequence. v_MHC1_ is the normalized bisecting vector of the parallelogram spanned by v_H1_ and v_H2_. Hence,

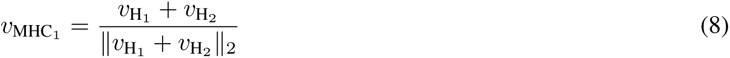

Let p_H1_ and p_H2_ be the center of mass estimated from C*_A_* atom coordinates of helices 1 and 2 respectively and v_H12_ the vector parallel to the line defined by points p_H1_, p_H2_ pointing towards helix 2. Then

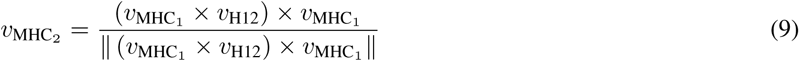

Where × is the cross product^2^. The centre of HLA mass p_MHC_ is defined as

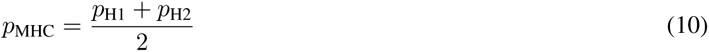

Using vectors v_MHC1_, v_MHC2_ and the HLA centre of mass p_MHC_ we define the HLA plane. The equation of the HLA plane for any given point k on the plane is given by:

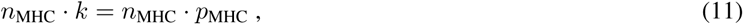

where

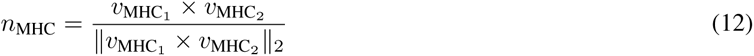

Given v*_rot_*,v_SB_, p_TCR_ for the TCR, along with v_MHC1_, v_MHC2_, p_MHC_ for the HLA and the HLA plane described by Eq 11 we can estimate the DoRIAT parameters. Visualization of v*_rot_*,v_SB_, p_TCR_, v_MHC1_, v_MHC2_ and p_MHC_ for the case of the 4FTV crystal structure can be found in Fig. 7. Let θ*^c^*, θ*^t^* and θ*^r^* be the cross (Rudolph et al., 2006), tilt(Teng et al., 1998) and roll(Teng et al., 1998) angles. Then

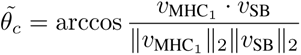

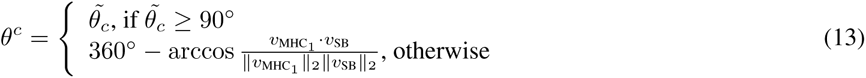

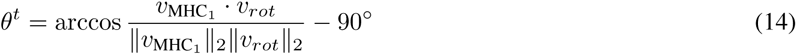

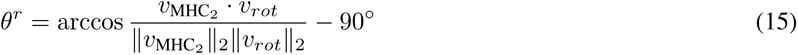

Let s*^z^* be the z axis displacement of p_TCR_ from the plane 11. then

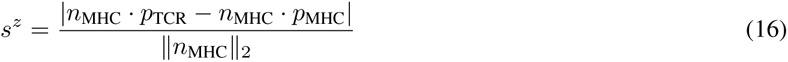

**Figure 7.**
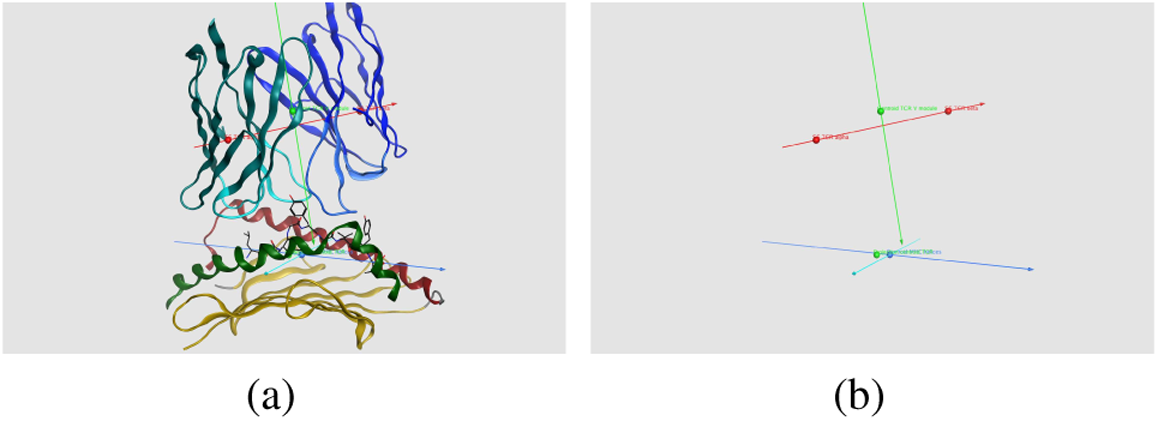
Vectors and points required to obtain binding mode parameters for 4FTV structure. Visual depiction of v_rot_(vector in light green), v_SG_(vector in red), v_MHC1_ (vector in blue), v_MHC2_ (vector in cyan), p_TCR_ (top light green sphere), p_MHC_ (blue sphere) with (Fig. 7a) and without (Fig. 7b) ribbon representation of the complex overlayed. The bottom light green sphere on the HLA plane is the projection of p_TCR_ used to estimate the displacements s^x^,s^y^ and s^z^.

**Algorithm 1.**
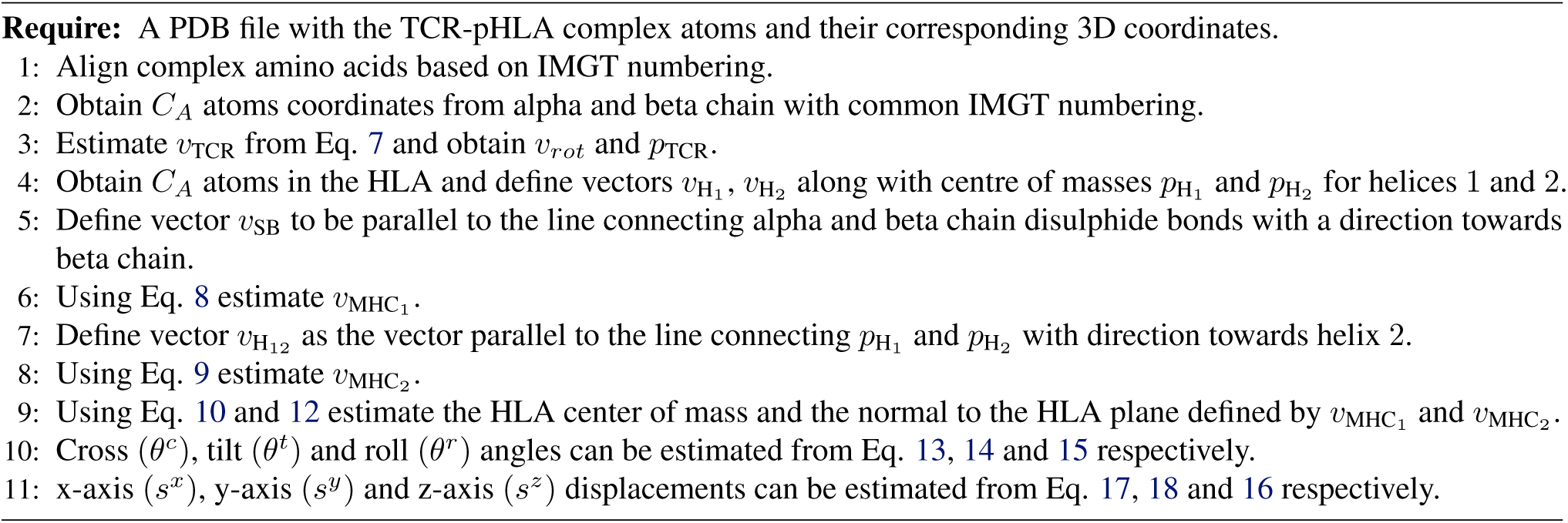
Pseudocode for estimating binding mode parameters.

Let s*^x^* and s*^y^* be the v_MHC_ and v_MHC_ axis displacements of p_TCR_ from p_MHC_ on the plane 11. Then

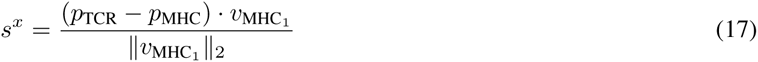

and

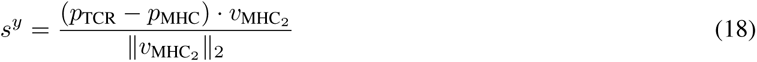

The procedure to estimate the binding mode parameters is summarized in Alg. 1/

### A.5. Data Preprocessing

The development of DoRIAT, involved a dataset of 58 crystal structures of human TCRs obtained from the Structural T-cell Receptor Database (STCRDab)(Leem et al., 2018)^3^ filtered to keep only ones that bind to HLA-A^∗^02. Using pdb-tools (Rodrigues et al., 2018) we remove all the non-protein atoms such as water, ions, other ligands for the TCR-pHLA complex.

Then alpha and beta chains were extracted into fasta format. Subsequently TCR alpha and beta chains are modeled using sequence-to-structure deep-learning based models.

Using MLDock we obtain 600 possible conformation of how the TCR-pHLA complex looks. From these docked structures, we calculate six parameters summarizing the global mode of TCR binding to the pHLA: cross, tilt, roll, shiftx, shifty, and shiftz. More information about these parameters can be found in Section A.4. To quantify the accuracy of the docked structures, we calculated the backbone distances from crystal using only C*_A_* atoms. Crystal structures with missing residues were ignored. Subsequently, backbone distances were normalized to a range between 0 and 1 by dividing with the largest observed distance plus 5 between the docked models and corresponding crystal structures. The normalized distance ranges were then converted to (-∞,∞) range to enable the seamless use of GP regressor from Tensorflow probability (Dillon et al., 2017). For model training and testing, we split the data based on complex id (TCR-pHLA) labels, maintaining a 3 to 1 ratio respectively.

As TCR-pHLA complexes contain four amino acid chains and the deep-learning methods examined are designed to score complexes containing only two chains, pdb files are preprocessed accordingly by merging alpha with beta chains and HLA with peptide in the described order.

### A.6. Implementation

For a training set of independent variables **X** = **x***_um_* u (1, …, U), (1, …, M) and dependent variables **Y** = **y***_um_* u (1, …, U), (1, …, M). To tune the GP kernel parameters l and ρ of Eq. 4 over training data we minimise the negative log marginal likelihood

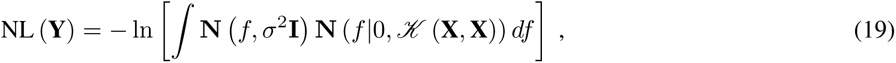

using Adam with learning rate 0.02. To minimise resource use, we have an early stopping policy applied if the negative log-likelihood drops by less than 1 unit in 10 epochs. To prevent over-fitting we tried cross validation but final parameters did not defer much.

### A.7. Mathematical Derivations

DoRIAT is trained and tested with response variable lying in the (-∞,∞) range. For predictions and model delineation dependent values have to be converted back into they original Å scale. In this section, we provide the detailed mathematical calculations required to avoid costly simulation along with the justification for using a non-standard approach to select the best model.

Suppose that DoRIAT for some input variable x^∗^ has predicted µ^∗^ and σ^∗^ as latent mean and standard deviation.

Let µ*_RMSD_* denote the RMSD average in observed space. To estimate µ*_RMSD_* we work as follows:

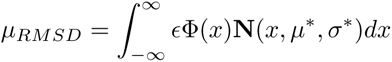

where Φ is the commulative cdf of standard normal and ɛ is the maximum observed RMSD we could observe. Using (Hartmann, 2017) we get the following result.

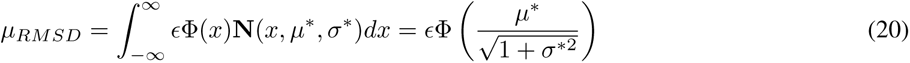

Let σ*_RMSD_* denote the RMSD standard deviation in observed space. To estimate σ*_RMSD_* we work as follows:

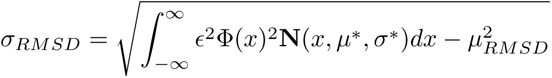

Again using (Hartmann, 2017) we get:

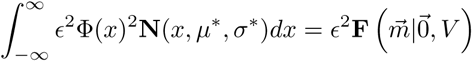

where **F** is the bivariate normal cdf with mean ^⃗^0 and variance V and

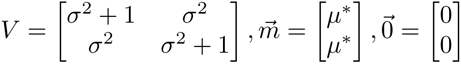

Hence:

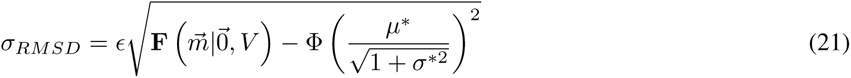

So, CV is estimated by:

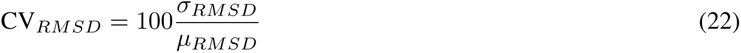

Predicted entropy E*_RMSD_*is given by the following formula:

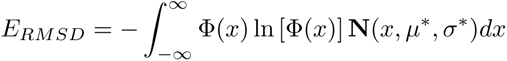

Since there is not closed form solution to the integral we first need to approximate −Φ(x) ln [Φ(x)]. A very good approximation can be obtained by Laplace approximation. By doing so we find that the mean is 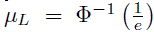 and standard deviation 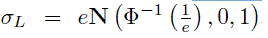. Using numerical integration on −Φ(x) ln [Φ(x)] we see that 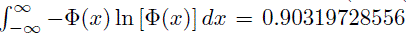. Hence c**N** (x, µ*_L_*, σ*_L_*) is a really good approximation of −Φ(x) ln [Φ(x)].

Hence:

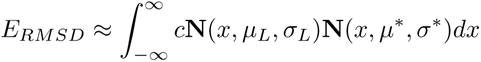

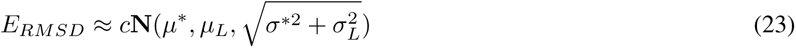

### A.8. Objective function justification

In this section we attempt to provide some additional insights as to why we chose Eq. 5 as our objective function.

To construct an objective function that selects good models we start by examining µ*_RMSD_*. Eq. 20 and Fig.8a indicate that if were to chose models based on µ^∗^ and σ^∗^, we would have a hard time as different combinations will yield the same results. Hence, it is important for our objective function to break such ties.

To resolve ties in model selection, we prioritize models with a high CV*_RMSD_*. In the face of uncertainty, we are making the optimistic assumption that we are overestimating µ*_RMSD_*. In practice we have observed that this assumption works well, as models close to the crystal structure present higher coefficient of variation compared to the remaining ones. By applying a constraint that retains the top 5% of models in terms of CV*_RMSD_*, we effectively filter out less favourable conformations. Then by selecting the model with the lowest µ*_RMSD_* among models within that subset we increase the chance of getting a desirable results.

**Figure 8.**
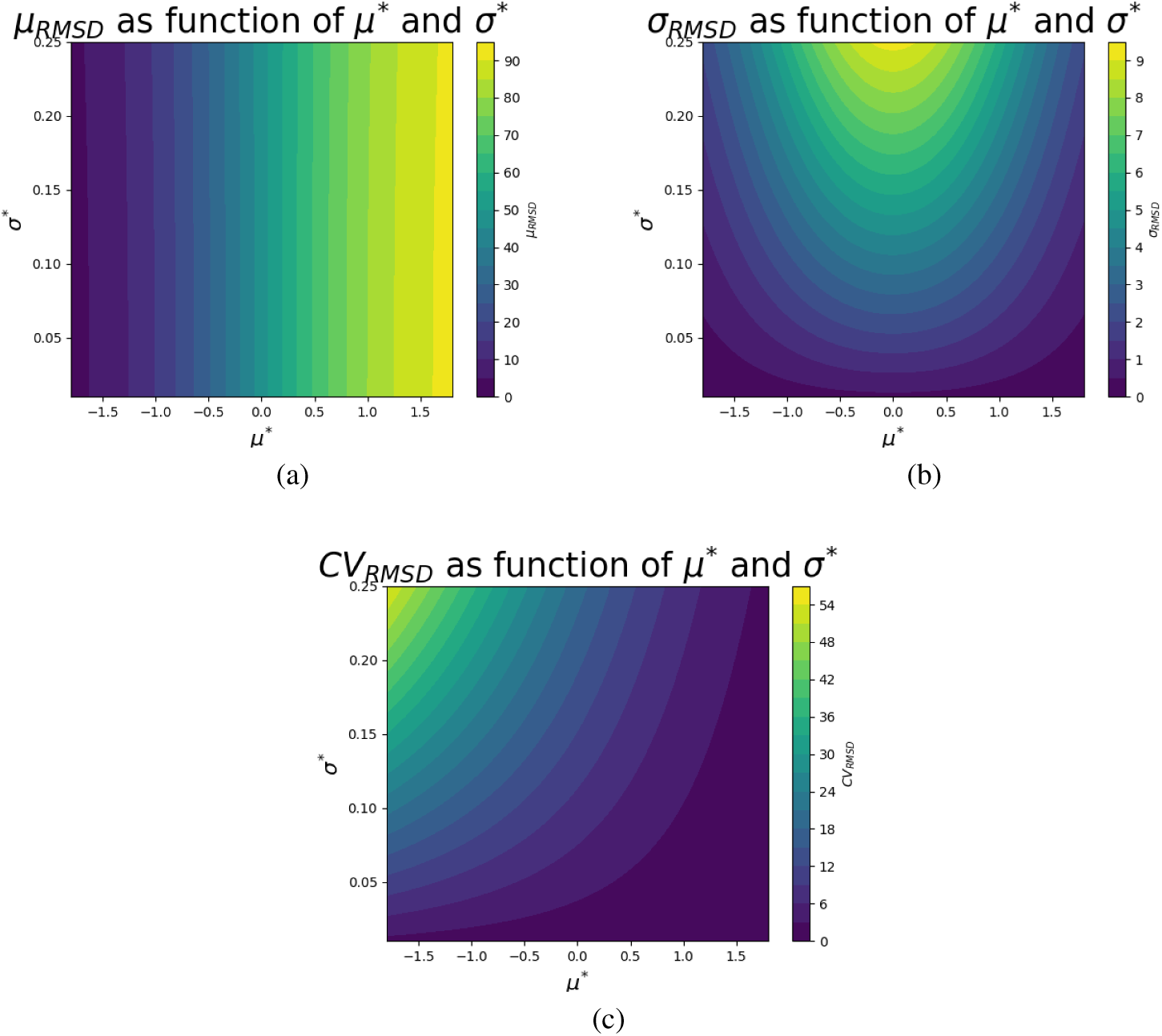
Contour plots of Eq. 20,21 and 22. Contour plots of predicted mean (µ_RMSD_, Fig 8a), standard-deviation (σ_RMSD_, Fig 8b) and coefficient of variation (CV_RMSD_, Fig 8c) as a function of parameters µ^∗^ and σ^∗^. Thresholds for µ^∗^ and σ^∗^ are estimated from test set observations.

### A.9. Cumulative hitrate across compared methods

In Sec. 3.2 we select a model from the docking output using different scoring approaches. Here we present a table summarizing the cumulative hitrate across compared methods.

**Table 5.** Table summarizing cumulative hitrate across compared methods. Table summarizing cumulative hitrate for DoRIAT, naive GNN, DeepRank-GNN, DeepRank-GNN-esm and MLDock across unseen complexes. Each method chooses one docked model that <u>believes it is closest to the crystal structure. These picks are then ranked using the measured RMSD</u> from crystal structure.

| Hitrate / Top | 1 | 10 | 20 | 40 | 70 | 120 | 170 | 250 |
| --- | --- | --- | --- | --- | --- | --- | --- | --- |
| DoRIAT | 22.22 | 77.78 | 77.78 | 88.89 | 94.44 | 100.00 | 100.00 | 100.00 |
| DeepRank<br>(naive-GNN) | 0.00 | 5.56 | 11.11 | 38.89 | 66.67 | 88.89 | 94.44 | 100.00 |
| DeepRank-GNN | 0.00 | 0.00 | 5.56 | 16.67 | 22.22 | 33.33 | 44.44 | 61.11 |
| DeepRank-GNN-esm | 0.00 | 5.56 | 5.56 | 11.11 | 16.67 | 27.78 | 33.33 | 55.56 |
| GNN-DOVE | 0.00 | 5.56 | 5.56 | 11.11 | 16.67 | 22.22 | 22.22 | 44.44 |
| MLDock | 0.00 | 11.11 | 11.11 | 16.67 | 38.89 | 50.00 | 72.22 | 88.89 |

### A.10. Benchmarking model selections with DockQ

In Sec. 3.2 we select a model from the docking output using different scoring approaches. Apart from ranking predictions based on RMSD, we also compare them using DockQ score. Since DockQ takes into account interface properties apart from ligand RMSD, it provides a more holistic assessment of the docking pose. Here we provide DockQ scores from the remaining test set and internal TCRs.

**Table 6:** Assessment of model selections using DockQ score for test set and internal TCRs. (Part2) Best perfroming selection is marked with bold. Cells are color-coded to indicate whether selection is canonical (green) or not (red) according to the thresholds of Tab. 2. DockQ scores are in the [0, 1] range.

|  | DoRIAT | DeepRank<br>(naive-GNN) | DeepRank-GNN | DeepRank-GNN-esm | GNN-DOVE | EMLy<sup>TM</sup>Dock |
| --- | --- | --- | --- | --- | --- | --- |
| TCR-pHLA complex C | 0.24 | 0.16 | 0.02 | 0.03 | 0.04 | **0.29** |
| 1QRN | **0.54** | 0.51 | 0.02 | 0.02 | 0.03 | 0.42 |
| 2BNQ | **0.20** | 0.09 | 0.01 | 0.02 | 0.04 | 0.06 |
| 2VLJ | **0.52** | 0.21 | 0.05 | 0.02 | 0.20 | 0.06 |
| 3QFJ | 0.51 | **0.60** | 0.10 | 0.58 | 0.56 | 0.59 |
| 4MNQ | 0.54 | **0.57** | 0.38 | 0.03 | 0.03 | 0.03 |
| 5C07 | **0.37** | 0.26 | 0.03 | 0.03 | 0.02 | 0.03 |
| 5COA | **0.32** | 0.1 | 0.11 | 0.02 | 0.02 | 0.05 |
| 5C0B | **0.41** | 0.19 | 0.03 | 0.02 | 0.03 | 0.04 |
| 5C0C | **0.25** | 0.09 | 0.03 | 0.03 | 0.03 | 0.03 |
| 5EU6 | **0.41** | 0.17 | 0.19 | 0.1 | 0.09 | 0.03 |
| 5MEN | **0.64** | 0.10 | 0.43 | 0.02 | 0.02 | 0.28 |
| 5YXU | **0.64** | 0.62 | 0.06 | 0.58 | 0.5 | 0.54 |

## Footnotes

1 A Python implementation of DoRIAT along with data pre-processing and docking data are available at https://zenodo.org/records/14763708

2 An alternative way of obtaining Eq. 9 is by starting with vectors *v*_MHC_ and *v*_H12_ and creating an orthonormal vector basis with Gram–Schmidt process (Datta, 2010).

3 Until November 2021.

